# Electricity-free nucleic acid extraction method from dried blood spots on filter paper for point-of-care diagnostics

**DOI:** 10.1101/2022.07.28.501845

**Authors:** Kenny Malpartida-Cardenas, Jake Baum, Aubrey Cunnington, Pantelis Georgiou, Jesus Rodriguez-Manzano

## Abstract

**Background:** Nucleic acid extraction is a crucial step for molecular biology applications, being a determinant for any diagnostic test procedure. Dried blood spots (DBS) have been used for decades for serology, drug monitoring, environmental investigations, and molecular studies. Nevertheless, nucleic acid extraction from DBS remains one of the main challenges to translate them to the point-of-care (POC).

**Method:** We have developed a fast nucleic acid extraction (NAE) method from DBS which is electricity-free and relies on cellulose filter papers (DBSFP). The performance of NAE was assessed with loop-mediated isothermal amplification (LAMP), targeting the human reference gene beta-actin. The developed method was evaluated against FTA cards and magnetic bead-based purification, using time-to-positive (min) for comparative analysis. We optimised and validated the developed method for elution (*eluted disk*) and disk directly in the reaction (*in-situ disk)*, RNA and DNA detection, and whole blood stored in anticoagulants (K_2_EDTA and lithium heparin). Furthermore, the compatibility of DBSFP with colourimetric detection was studied to show the transferability to the POC.

**Results:** The proposed DBSFP is based on grade 3 filter paper pre-treated with 8% (v/v) igepal surfactant, 1 min washing step with PBS 1X and elution in TE 1X buffer after 5 min incubation at room temperature, enabling NAE under 7 min. Obtained results were comparable to gold standard methods across tested matrices, targets and experimental conditions, demonstrating the versatility of the methodology. Lastly, *eluted disk* colourimetric detection was achieved with a sample-to-result turnaround time under 35 min.

**Conclusions:** The developed method is a fast, electricity-free, and low-cost solution for NAE from DBSFP enabling molecular testing in virtually any POC setting.

## Introduction

Dried blood spot (DBS) technology has been widely used for sample collection and diagnostic purposes^1–3^. One of the most significant advantages of DBS is that whole blood can be safely stored and shipped at room temperature (RT) after simply spotting a few microliters onto cellulose filter paper (DBSFP).^4–6^ Several DBSFPs are commercially available for the collection of whole blood such as FTA classic cards, Whatman 903 protein saver cards or most recently, FTA elute cards (QIAGEN, Whatman, Cytiva); all of them allowing for cell lysis and the preservation of the genetic material for a long period of time without the need for cold chain storage. However, they are expensive (£3-5 per single card) and commonly treated with non-disclosed proprietary compounds (Table S1). Alternatively, filter papers which have not been specifically designed for blood collection have been evaluated for nucleic acid extraction (NAE), such as Fusion 5, filter paper grade 1 or filter paper grade 3, demonstrating their potential for nucleic acid detection for clinical purposes.^5,7^ Nevertheless, current strategies for NAE from DBSFP still rely on: (i) silica-based spin columns (QIAamp DNA Blood Mini Kit, Maxwell RSC DNA FFPE Kit or ABI 6100 Nucleic Acid Prep Station protocol)^8,9^, (ii) magnetic beads^10^, (iii) long incubations at high temperatures (95 °C)^11,12^, or (iv) long protocols with hazardous chemicals^13,14^. Although, simplification of these methods has been developed, they have only been applied for their direct use in PCR^7,15^, limiting downstream analysis to laboratory-based settings. The high demand of rapid testing at the point-of-care (POC) is creating a need for the development of NAE method from DBSFPs that must be: (i) compatible with isothermal chemistries such as loop-mediated isothermal amplification (LAMP) to reduce the hardware complexity needed to perform nucleic acid amplification^16^, (ii) independent of laboratory equipment to be accessible in limited-resource settings, (iii) affordable and compatible with broadly available molecular reagents to reduce the cost and increase scale-up capabilities, and (iv) stable at RT conditions to avoid the use of cold-chain storage.^17^

In this work, addressing aforementioned requirements, we report a fast (under 7 min) NAE method from DBSFP which is electricity-free, performed at RT and relies on cellulose filter paper pre-treated with the surfactant igepal. The performance of the method was assessed with LAMP targeting the human gene beta-actin, and optimised for elution (*eluted disk*) and disk in the reaction (*in-situ disk)*. The developed method allows for RNA/DNA purification and was compatible with whole blood stored in anticoagulants. In addition, end-point colorimetric detection of amplified products was performed to show its applicability to the POC, showing a sample-to-result turnaround time under 35 min. The experimental design and optimised NAE protocol is depicted in Figure 1.

**Figure 1.**
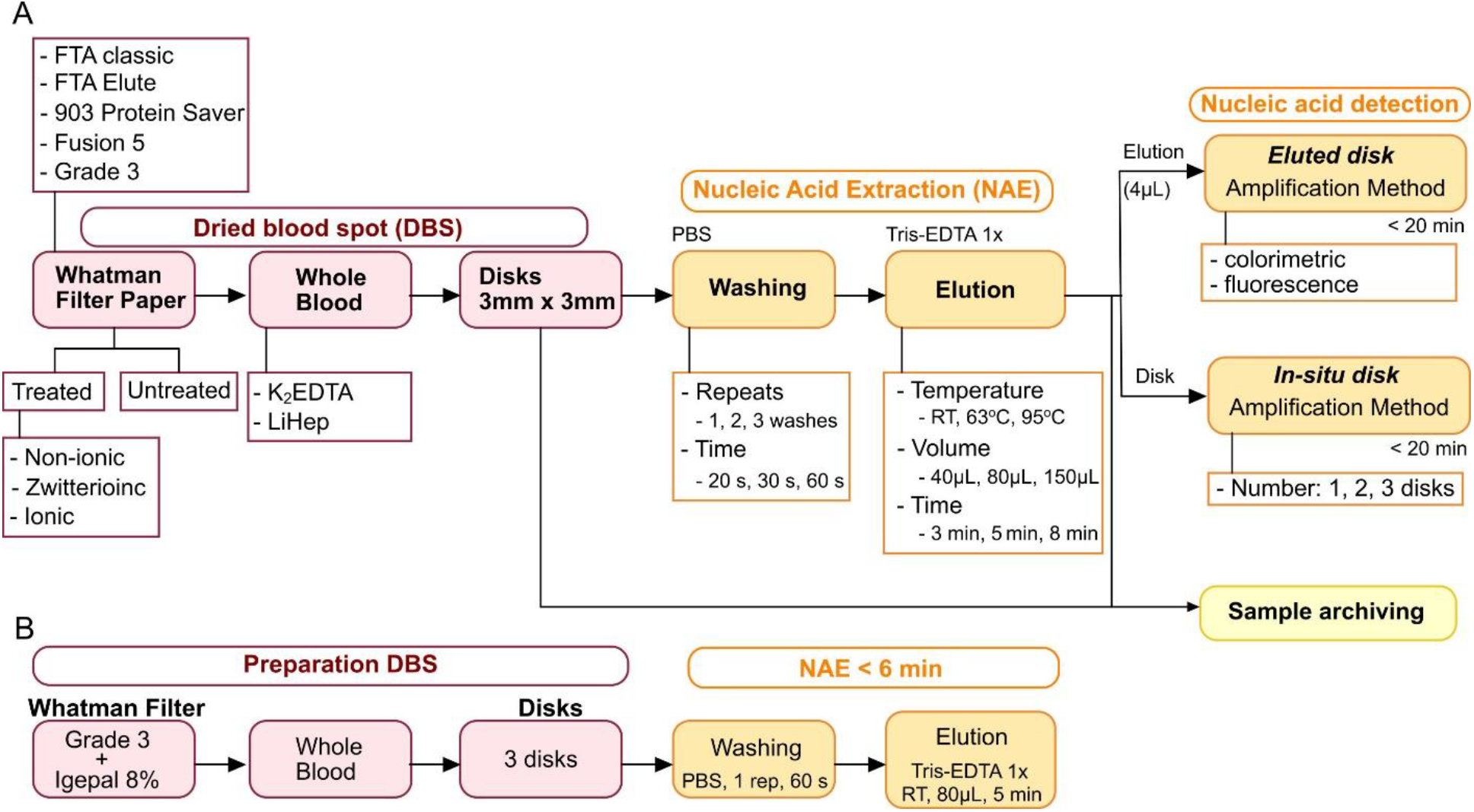
Experimental design used in this study. (A) Workflow for optimisation of NAE from DBSFP. (B) Optimised DBSFP workflow.

## Methods

### Samples and controls

Whole blood samples stored in K_2_EDTA or lithium heparin (LiHep) were purchased from Cambridge Bioscience (BLD1DA2EDT10-XS, BLD1DC4LHE10-FS) and kept at 4 °C until further use. Then, it was dried onto filter papers by pipetting 2 µL each time to standardise the collection process, and compare the different filter papers without the influence of their physico-chemical properties at the time of generating the DBSFP. Human Genomic DNA (G1521, Promega) was used as positive control and synthetic DNA was used to validate the LAMP assay performance. A gBlock was purchased from Integrated DNA Technologies (IDT), resuspended in 1× TE buffer and the concentration was quantified (copies per µL) using a Qubit fluorometer instrument. Sequence of the gBlock is shown in Table S2.

### LAMP primer design

A LAMP assay targeting the human house-keeping gene beta-actin (ACTB) was designed using the software Primer Explorer version 5.0 (http://primerexplorer.jp/lampv5e/index.html), based on the sequences retrieved from NCBI GenBank database (https://www.ncbi.nlm.nih.gov/genbank/) with accession number: NC_000007.14:c5530601-5527148, and NM_001101.5. Sequences were aligned using the MUSCLE algorithm^18^ in Geneious 10.0.5 software^19^, and primers were designed to detect both mRNA and DNA. All primers used in this study were purchased from IDT and resuspended in TE 1X buffer at 200 µM. Primer sequences are provided in Figure 2C.

**Figure 2.**
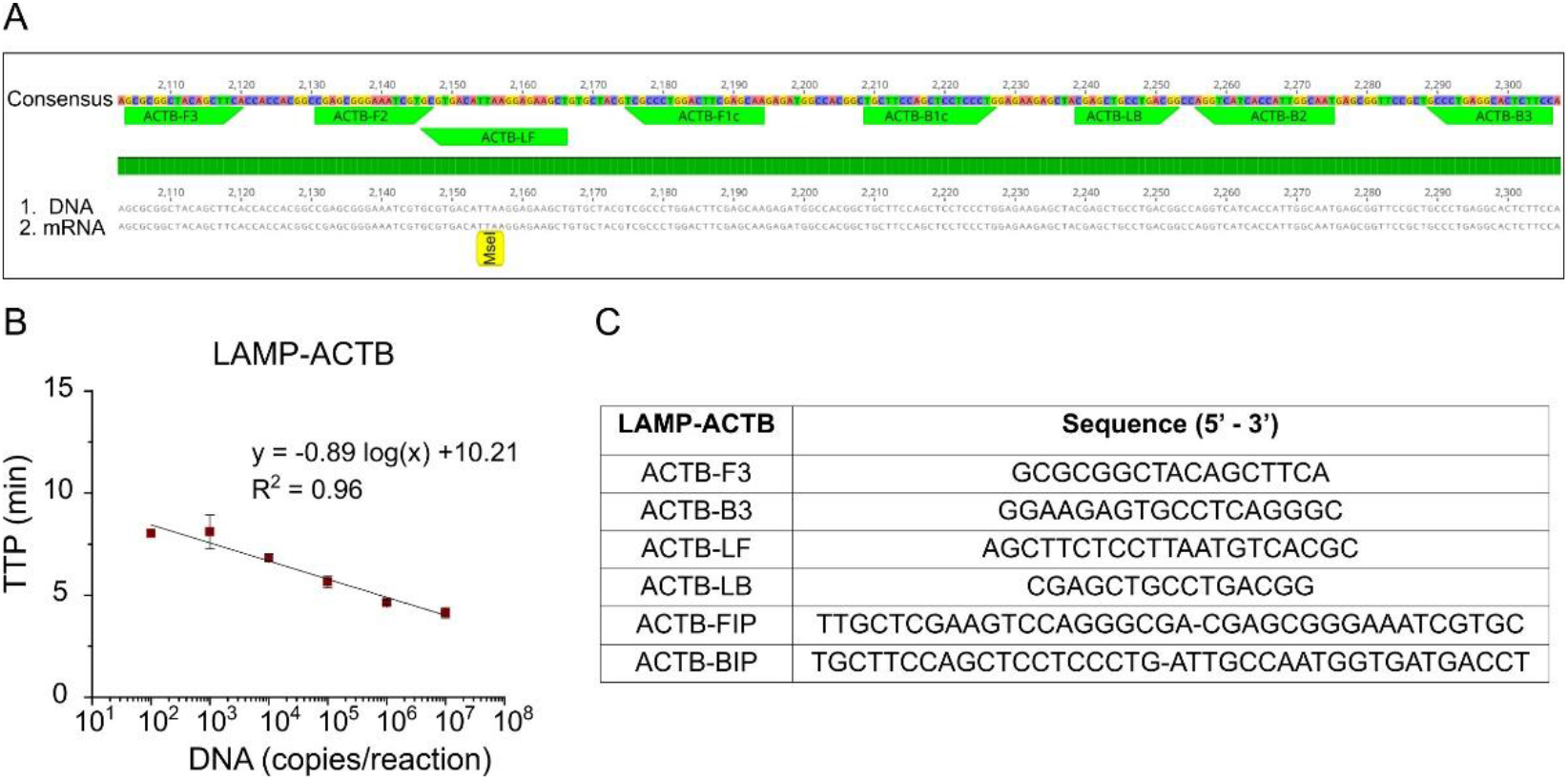
LAMP assay for the detection of the human house-keeping gene beta-actin. (A) DNA and mRNA alignment showing the primer regions. (B) Standard curve using the LAMP-ACTB assay with synthetic DNA dilutions ranging from 10 to 10^7^ copies per reaction. (C) LAMP-ACTB primer sequences.

### LAMP reaction conditions

(i) Fluorescent detection: LAMP reactions were carried out at a final volume of 10 µL per reaction. Each mix contained the following: 1 µL of 10× custom isothermal buffer (pH 8.5 – 9), 0.5 µL of MgSO4 (100 mM stock), 0.56 µL of dNTPs (25 mM stock), 0.6 µL of BSA (20 mg/mL stock), 1 µL of 20× LAMP primer mix (F3/B3 5 µM, LF/LB 20 µM and FIP/BIP 40 µM), 0.25 µL of Syto9 dye (20 µM stock), 0.25 µL of NaOH (0.2 M stock), 0.04 µL of Bst 2.0 WarmStart DNA polymerase (120K U/mL stock), 0.3 µL of WarmStart RTx Reverse Transcriptase (15K U/mL stock), 4 µL sample, and enough nuclease-free water (Invitrogen) to bring the volume to 10 µL. In the case of the *eluted disk* method, 4 µL of elution from DBSFP and enough nuclease-free water (Invitrogen) were used to bring the final volume to 10 µL. For the *in-situ disk* method, reactions were carried out at a final volume of 20 µL (2 × 10 µL reaction) adding enough nuclease-free water and immersing the disk(s) into the reaction; disk(s) still wet after the incubation step in the NAE from DBSFP. Reactions were loaded into 96-well plates and were performed at 63 °C during 35 min using a LightCycler 96 Real-Time System (LC96) (Roche Diagnostics). One melting cycle was performed at 0.1 °C/s from 63 °C up to 97 °C for validation of the specificity of the amplified products. Non-template control (NTC) was included in every experiment.

(ii) Colorimetric detection: LAMP reactions were performed at a final volume of 20 µL using the WarmStart Colorimetric LAMP 2× Master Mix (10 µL), 20× LAMP primer mix (2 µL), 4 µL of the elution and enough nuclease-free water to reach a final volume of 20 µL. Amplification reactions were carried out in a portable thermal block (N2400-4020 and N2400-4021, Starlab) during 35 min at 63 °C (checking at 20 min).

### Analytical sensitivity of the LAMP-ACTB assay

Analytical sensitivity was evaluated using 10-fold dilutions of synthetic DNA in nuclease-free water ranging from 10^1^ to 10^7^ copies per reaction. Each condition was run in triplicates using the LC96 real-time instrument.

### Analytical specificity of the LAMP-ACTB assay

Analytical specificity of the amplified products was evaluated by melting curve analysis. To further demonstrate the specificity among the proposed *eluted disk* and *in-situ* disk methods, restriction digestion of the LAMP products was performed using the *MseI* enzyme (NEB, UK) which performs a single cut at T/TAA position. The restriction was performed for 15 min at 37 °C, using the rCutSmart buffer provided by the manufacturer. Enzyme was deactivated following manufacturer’s protocol by incubating for 20 min at 65 °C. The digested products were analysed by electrophoresis on a 1.5% agarose gel stained with SYBR Safe DNA Gel Stain (InvitroGen). The experiment was run for 1 h at 145 V and the gel was visualized under UV light using a BioSpectrum Imaging System (Ultra-Violet Products Ltd.). The Quick-Load 1 kb Plus DNA Ladder (NEB, UK) was used as reference.

### Protocol for custom pre-treatment of filter papers

Detergents used in this study were purchased from Sigma-Aldrich and included: sodium dodecyl sulfate (L3771-25G), CHAPS (C9426-1G), Triton X-100 (T8787-100ML), Tween 20 (P9416-50ML) and Igepal (I8896-50ML). Detergents were diluted in nuclease-free water at final concentrations of 2%, 4%, 6%, 8% and 10% w/v or v/v. A total of 150 µL of each solution was deposited on the assessed filter papers and were left to dry overnight at RT.

### Experimental conditions used to optimise the NAE from DBSFP

The NAEs were performed as follows: (i) whole blood was spotted on the pre-treated or untreated filter paper and was left to dry overnight (following standard practice^20^); (ii) 1, 2 or 3 disks of 3mm × 3mm each were punched using the Harris Core puncher from the UniCore Punch Kit 3.00 mm; disks were deposited into a 1.5 mL or 2.0 mL Eppendorf tube and the puncher was rinsed with 70% ethanol after every use; (iii) disk(s) washing was performed with PBS 1X by shaking, vortex or passively (not disturbed). The three methods were assessed at varying conditions including number of washes (1, 2 or 3), time per wash (20 s, 30 s or 60 s), and total volume of PBS 1X (400 µL, 600 µL, or 1500 µL); (iv) nucleic acid elution was performed by adding TE 1X buffer at different volumes (40 µL, 80 µL, or 150 µL), different incubation times (3, 5 or 8 min) at varying temperatures (RT, 63 °C, or 95 °C); (v) Lastly, for the *eluted disk* method, 4 µL of the elution was added into the LAMP reaction (final volume of 10 µL), and for the *in-situ disk* method, 1, 2 or 3 disks were added to a LAMP reaction with final volume of 20 µL to completely cover the disk in solution (disks still being wet when transferred).

### Magnetic bead-beads nucleic acid extraction from whole blood

Magnetic beads-based extraction was performed using the Dynabeads DNA DIRECT Blood kit from Invitrogen (Life Technolgies) following the manufacturer’s instructions. Input sample volume was 100 µL.

### Statistical analysis

Time-to-positive (TTP) data is presented as mean TTP ± standard deviation; p-values were calculated by Welch’s unequal variance two sample t-test or one-way ANOVA with Fisher LSD means comparison in Origin 2019 (v9.6). A p-value equal to 0.05 was considered as the threshold for statistical significance.

## Results

### LAMP assay for detection of the house-keeping gene beta-actin

A novel LAMP assay, named LAMP-ACTB, was designed targeting the human house-keeping gene ACTB. Based on the genomic sequences NC_000007.14:c5530601-5527148 and NM_001101, the assay was optimised to detect both mRNA and DNA (Figure 2A). Primer sequences are shown in Figure 2C. The ACTB gene is highly expressed, particularly in blood, and widely used as reference gene.^21,22^ Analytical sensitivity was assessed using synthetic DNA, achieving a limit of detection of 100 copies per reaction within 10 min. The amplification standard curve is shown in Figure 2B, with a coefficient of determination of R^2^ = 0.96. NTCs were negative during the 35 cycles run, and non-specific products were not observed. The lowest tested concentration (10 copies per reaction) did not amplify.

### Evaluation of different detergents for enhanced dried blood spot lysis

Detergents with different physico-chemical properties were selected to evaluate their lysis effect on DBSFP, as they are the major ingredient that determines the lysis strength of a given lysis buffer.^23^ Detergents used in this study include: (i) ionic such as sodium dodecyl sulfate (SDS); (ii) zwitterionic such as CHAPS; and (iii) non-ionic such as Triton X-100, Tween 20 and Igepal (Nonidet-40 type). Properties and applications of these detergents are summarised in Table S3.

The detergents were dried onto FTA classic cards at different concentrations ranging from 0% to 10% v/v or w/v, where 0% denoted the absence of detergent. Nucleic acid extraction was performed from whole blood in K_2_EDTA using the *eluted disk* method and including a washing step (3 washes of 20 sec with 400 µL pf PBS 1X) and incubation at RT, 63 °C or 95 °C during 8 min. Elutions were tested with the LAMP-ACTB assay targeting both DNA and RNA of the house-keeping gene ACTB. Results are shown in Figure 3, where the detergents at different concentrations are plotted against TTP values. Nucleic acid amplification was comparable between incubation at RT and at 95 °C (p-value > 0.05), all with TTP values below 15 min. Results obtained at 63 °C were inconsistent and negative for most of the conditions (data not shown).

**Figure 3.**
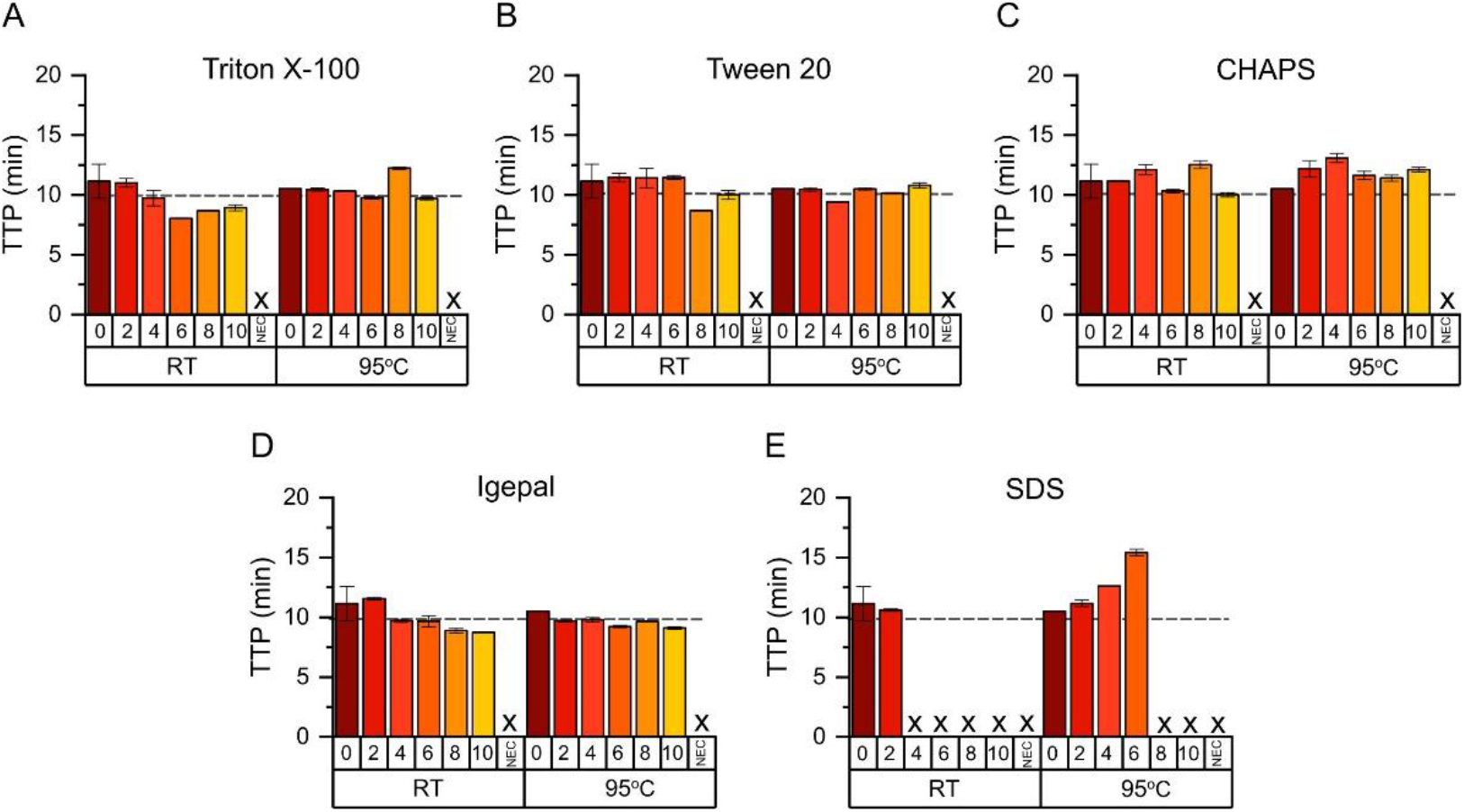
Evaluation of detergents for DBSFP lysis with the *eluted disk* method. Detergent dilutions ranging from 2% to 10% v/v or w/v were spotted in the sample area of FTA classic cards. After blood was dried on them, NAE was performed by washing and incubating at RT or 95 °C. TTP values from the amplification of the human ACTB gene using the LAMP-ACTB assay are plotted against the different detergent concentrations and incubation temperatures. (A) Triton X-100, (B) Tween 20, (C) CHAPS, (D) Igepal and (E) SDS. The “0” denotes no detergent addition and NEC denotes negative extraction control which consisted of disks without dried blood; for each condition, n = 3.

The effect of the detergents to enhance cell lysis was demonstrated by the lower TTP values obtained at RT conditions as the detergent concentration was increased. The high strength of the ionic detergent SDS resulted inhibitory at concentrations above 2%. Increasing the percentage of detergent did not improve lysis performance when pre-incubating at 95 °C. Thermal lysis is commonly achieved at high temperatures such as 95 °C, being the addition of detergent negligible. Among the different detergents, igepal showed a linear trend as the concentration was increased with a coefficient of determination of R^2^ = 0.85 (Figure S1). Deposition of igepal at 8% (p-value = 0.04, compared to absence of detergent) prior to blood collection was the condition selected for subsequent experiments.

### Comparison of filter papers for nucleic acid extraction

Filter papers with different physico-chemical properties were evaluated for their blood collection capability and NAE performance. Filter papers used in this work included: FTA classic cards, Grade 3 Qualitative Filter Paper Standard Grade, Fusion 5, 903 Protein Saver cards, and FTA elute cards. Photographs of the dried blood spots are shown in Figure 4A, and properties of the filter papers are summarised in Table S1. Comparison between the recommended protocol for FTA elute cards (30 min incubation at 95 °C) and the *eluted disk* method (8 min incubation at 95 °C) is shown in Figure S2 to demonstrate that the benefit of reducing the incubation time did not have a negative effect in the NAE step.

**Figure 4.**
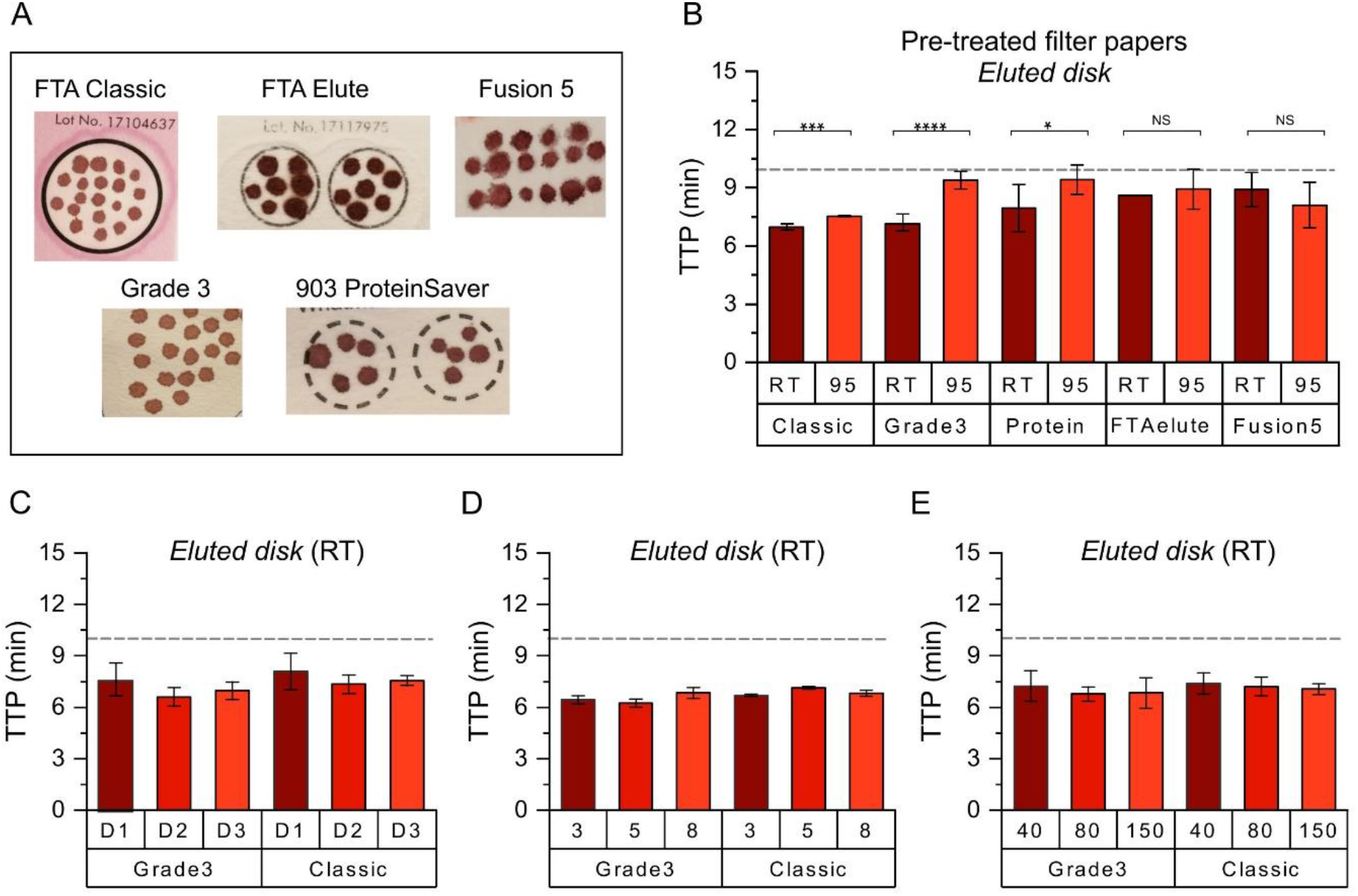
Optimisation of NAE from DBSFP with *eluted disk* method. (A) Photographs of DBSFP after the pre-treatment with igepal 8% v/v: FTA classic cards (Classic), FTA elute card (FTAelute), Fusion 5, grade 3 qualitative filter paper (Grade3) and whatman 903 Protein Saver cards (Prtoein). (B) Elutions from incubation at RT or 95 °C using different filter papers. (C) Elutions from incubation at RT using one disk (D1), two disks (D2) or three disks (D3) from grade 3 filter paper or FTA classic cards. (D) Elutions from incubation at RT using three disks during different times: 3 min (3), 5 min (5) or 8 min (8) from Grade3 or Classic filter papers. (E) Elutions from incubation at RT using three disks during 5 min eluted at different volumes of TE 1X buffer: 40 µL (40), 80 µL (80) or 150 µL (150) using grade 3 filter paper or FTA classic cards. From (B) to (E), TTP values from the amplification of the human ACTB gene using the LAMP-ACTB assay. Dash line at y-axis sets a threshold at 10 min; for each condition, n = 4.

Untreated and pre-treated filter papers with igepal at 8% (I8) were evaluated. NAE was performed from whole blood in K_2_EDTA using the *eluted disk* method and including a washing step (3 washes of 20 sec with 400 µL pf PBS 1X) and incubation at RT, 63 °C or 95 °C during 8 min. Elutions were tested with the LAMP-ACTB assay. Results in Figure 4B showed that RT incubation resulted in faster nucleic acid amplification compared to 95 °C for Classic (p-value < 0.001), Grade3 (p-value < 0.0001) and Protein filter papers (p-value < 0.05). Furthermore, results obtained with the Classic and Grade3 filter papers after incubation at RT were significantly different from the other filter papers (p-value < 0.05), with TTP values of 6.90 ± 0.16 min and 6.87 ± 0.43 min, respectively. Results obtained with the elutions from incubation at 63 °C where inconsistent and mostly negative (data not shown). Untreated filter papers showed a similar trend to pre-treated filter papers, but only Grade3 showed significant lower TTP at RT compared to 95 °C (Figure S2, p-value < 0.0001). The absence of detergent resulted in higher TTP values at RT and 95 °C for every filter paper except for FTA elute cards due to the chemical properties of this type of cards, being the addition of detergent negligible.

### Optimisation of conditions for nucleic acid extraction

Optimisation of nucleic acid extraction from DBSFP using whole blood in K_2_EDTA and the *eluted disk* method was carried out using pre-treated Classic and Grade3 filter papers to evaluate the effect of: (i) number of disks; (ii) pre-incubation time; (iii) volume of the elution; and (iv) elution buffer. Incubation temperatures at RT, 63 °C and 95 °C were tested across all the conditions, except for (iii) and (iv) which were only tested at RT.

Firstly, one, two and three disks (D1, D2 and D3, respectively) were washed (3 washes of 20 sec with 400 µL pf PBS 1X) and incubated. A higher number of disks was correlated to a lower TTP value (Figure 4C). Secondly, different incubation times of 3, 5 and 8 minutes were evaluated with three disks. Results in Figure 4D showed that TTP values were comparable across the three pre-incubation times at RT conditions, being both Classic and Grade3 filter papers performing with a similar trend. Elutions from incubation at 95 °C were comparable to RT conditions (Figure S3), with RT incubation resulting in faster and more consistent nucleic acid amplification. Incubation at 63 °C showed inconsistent results and no amplification was observed in most of the cases (data not shown). Lastly, elution buffers and volumes were evaluated with 3 disks and incubated during 5 min at RT (selected as the most optimal conditions). Results obtained showed no significant difference across the volumes tested, but lower standard deviation was obtained with 80 µL (Figure 4E). Elution in TE buffer 1X resulted in faster nucleic acid amplification compared to elution in nuclease-free water (Figure S3) due to the higher pH of the TE buffer and its chemical properties to preserve nucleic acids. Across all the parameters assessed for the optimisation of nucleic acid extraction in a rapid and efficient way, an incubation time of 5 min at RT in 80 µL of TE and the use of 3 disks were selected.

### Duality of the DBSFP

The NAE method, using whole blood in K_2_EDTA, was further assessed by directly placing the disks into the LAMP reaction, named *in-situ disk* method. Firstly, *in-situ disk* method was evaluated at different conditions including number of disks, incubation time, and elution volume. No significant differences were observed among them as shown in Figures 5A-C, respectively. Secondly, the washing step was evaluated at various conditions, including: (i) no wash (W1), (ii) the use of a vortex (W2 to W4), (iii) shaking (W5 and W6), (iv) and passive washing (W7 and W8). Among them, lower TTP values were obtained with W4 and W5 which consisted of 3 washes using vortex and 1 wash by shaking, respectively. Between them, W5 was chosen for subsequent experiments (Figure 5D). Thirdly, disks incubated at the three different temperatures (RT, 63 °C and 95 °C) were tested. Contrary to the *eluted disk* method, nucleic acid amplification was detected with the disks after incubation at the three conditions. TTP values were significantly lower at RT and 95 °C compared to 63 °C (Figure 5E). Average TTP values were 5.00 ± 0.51 min, 10.45 ± 1.15 min and 5.27 ± 0.96 min for RT, 63 °C, and 95 °C, respectively. As reference, the absence of an incubation step was also evaluated. Results showed a higher variability, and TTP values were significantly higher (8.71 ± 2.65 min) than those obtained at RT or 95 °C (Figure 5E). Also, the *in-situ disk* method was performed with the different filter papers and results (Figure S4) agreed with those obtained in Figure 5E. Finally, in order validate the specificity of the obtained amplification products, restriction analysis was performed using the *MseI* restriction enzyme. As shown in Figure 5F, a band at 152 bp and 51 bp could be observed demonstrating the specific amplification of the LAMP-ACTB assay with both, the *eluted disk* and *in-situ disk* methods.

**Figure 5.**
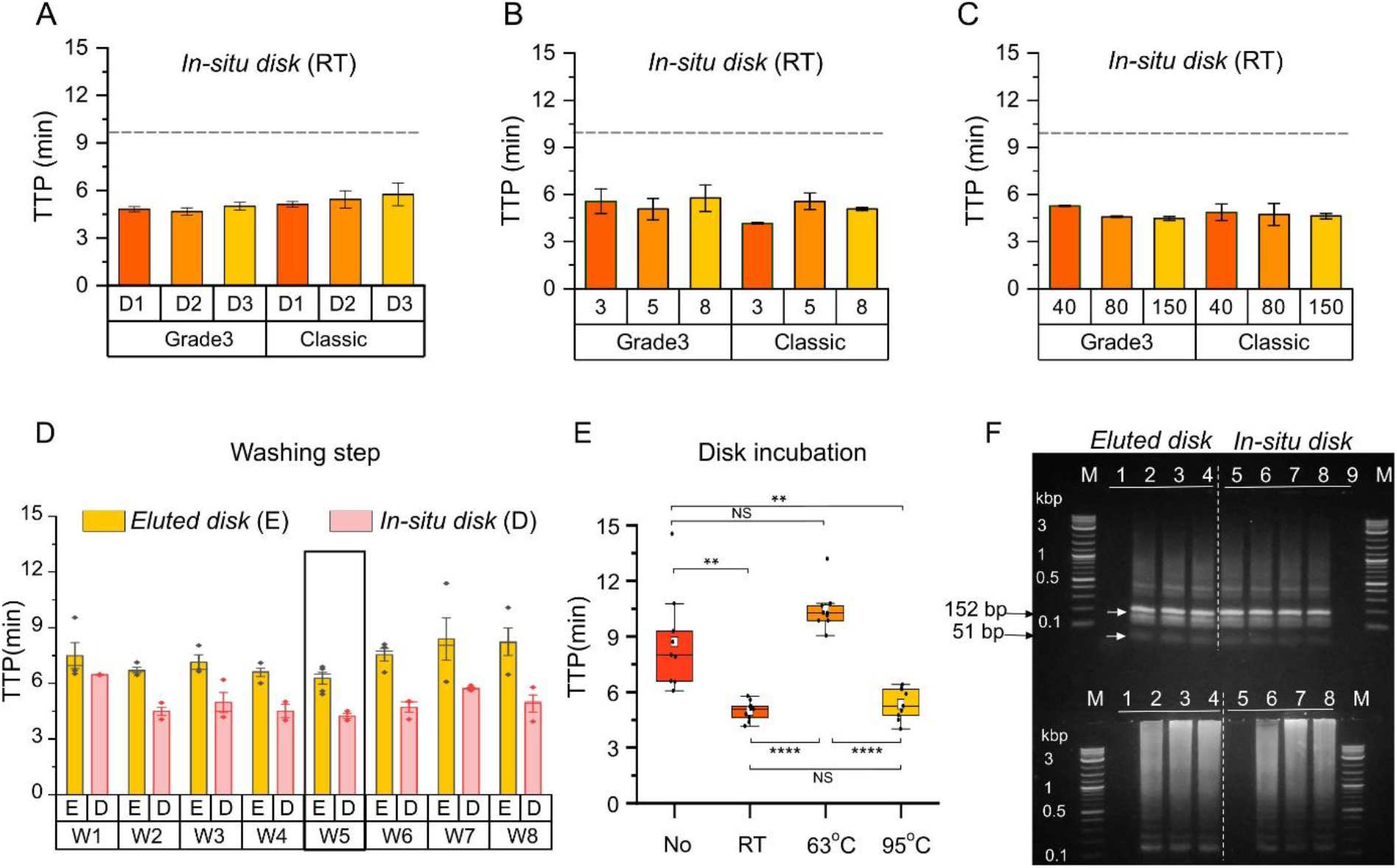
Duality of the DBSFP. (A) TTP values from LAMP amplification after *in-situ disk* method with RT incubation using one disk (D1), two disks (D2) or three disks (D3) from Grade3 and Classic cards. (B) TTP values from LAMP amplification after *in-situ disk* method with RT incubation during different times: 3 min (3), 5 min (5) or 8 min (8) using three disks from Grade3 and Classic cards. (C) TTP values from LAMP amplification after *in-situ disk* method with RT incubation during 5 min eluted at different volumes: 40 µL (40), 80 µL (80) or 150 µL (150) using three disks from Grade3 and Classic cards. (D) Optimisation of the washing step for *eluted disk* and *in-situ disk* methods. W1 (no wash) W2 (1 wash, 1500 µL); W3 (2 washes, 600 µL); W4 (3 washes, 400 µL); W5 (1 wash, 1500 µL); W6 (2 washes, 600 µL); W7 (1 wash, 1500 µL); W8 (2 washes, 600 µL); W2 to W4 using a vortex, W5 and W6 by shaking, and W7 and W8 passive. The washing step was performed during 1 min overall. (E) Boxplot showing the effect of the incubation step and its temperature for the *in-situ disk* method. “No” denotes the absence of incubation step after washing. Each dot represents a sample, n = 9; horizontal lines in the boxes indicate medians; lower and upper edges of boxes indicate interquartile range and whiskers are <1 times the interquartile range. (F) Restriction analysis and gel electrophoresis of amplified products. Specificity of the LAMP-ACTB assay and NAE methods is confirmed in the top part of the gel after digestion with the RE *MseI* showing bands at 152 bp and 51 bp. Lane M shows Quick-Load 1 kb Plus DNA Ladder, lane 1 and 9 show NTC, lane 2-4 show digested amplified products from *eluted disk* method at RT, 63 °C and 95 °C respectively; lane 5-9 show digested amplified products *in-situ disk* method at RT, RT, 63 °C and 95 °C respectively. Low part of the gel shows amplified products, not digested. Lane M shows Quick-Load 1 kb Plus DNA Ladder, lane 1 and 5 show NTC, lane 2-4 show amplified products from *eluted disk* method at RT, 63 °C and 95 °C respectively; lane 6-8 show amplified products from *in-situ disk* method at RT, 63 °C and 95 °C respectively.

### Validation with samples and detection of RNA and DNA

Whole blood samples stored in anticoagulants were screened for RNA and DNA purification from DBSFP with the *eluted disk* method (Figure 6A). Elutions from whole blood stored in K_2_EDTA spotted on Grade3 filter paper reported an average TTP value of 6.99 ± 0.45 min and 18.76 ± 1.34 min with and without the addition of the reverse transcriptase enzyme, respectively. Elutions from whole blood stored in LiHep reported TTP values of 7.62 ± 0.60 min and 7.50 ± 0.63 min with and without the reverse transcriptase with the Grade3 filter paper.

**Figure 6.**
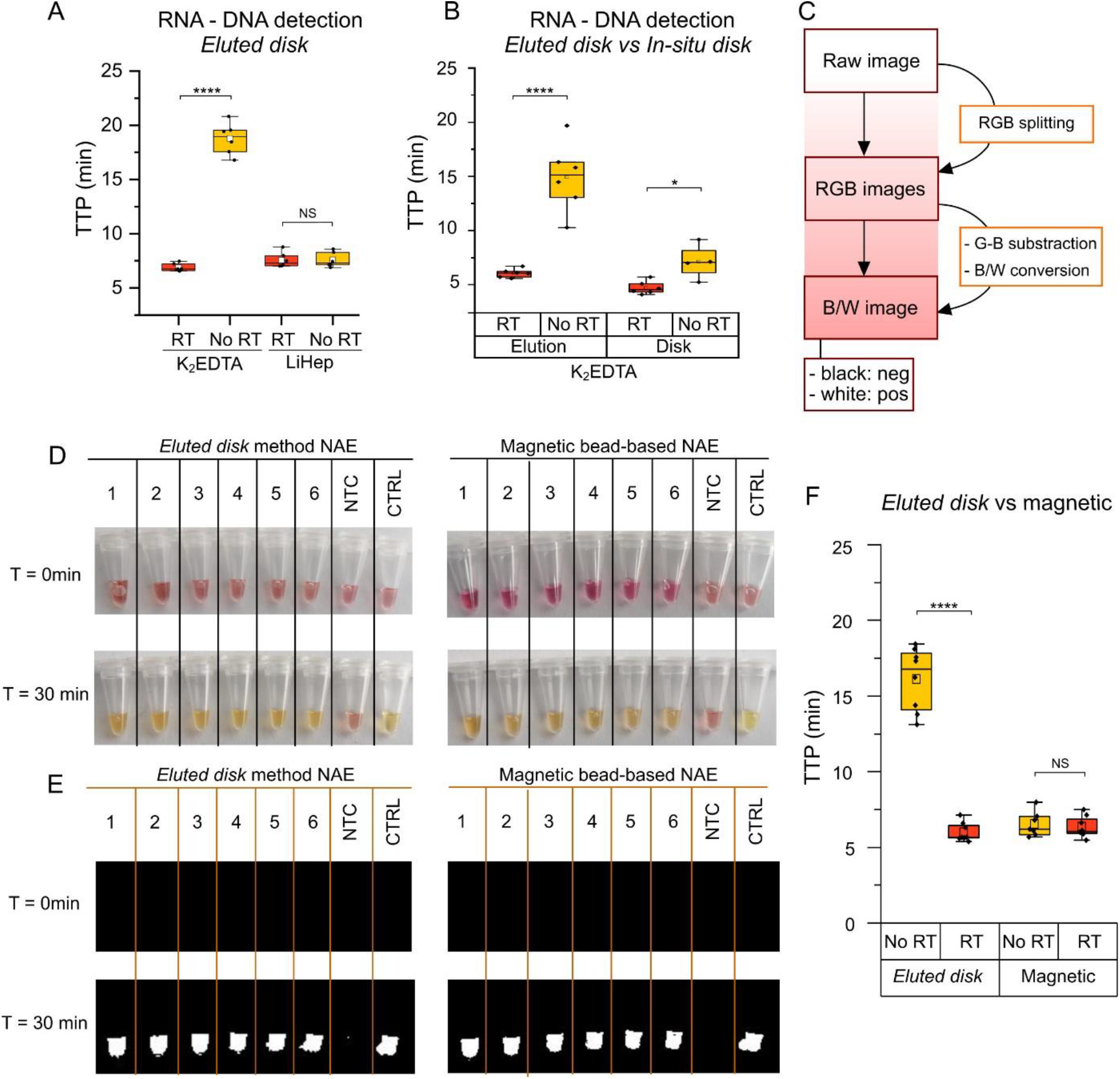
Detection of DNA/RNA using colorimetric LAMP. (A) Boxplot showing RNA and DNA detection from whole blood stored in anticoagulants using the *eluted disk* method. (B) Boxplot showing RNA and DNA detection from whole blood in K_2_EDTA using the *eluted disk* and *in-situ disk* method. Each dot represents a sample, n = 6; horizontal lines in the boxes indicate medians; lower and upper edges of boxes indicate interquartile range and whiskers are < 1 times the interquartile range. (C) Workflow for colorimetric image processing. Algorithm implemented in MATLAB from raw image to a binary output. (D) Colorimetric diagnostics from sample-to-result with the *eluted disk* method from DBSFP (left) and a commercial magnetic bead-based NAE method (right). Samples used included whole blood in K_2_EDTA (1-3) and whole blood in LiHep (4-6); non-template control (NTC); and positive control (CTRL), purified human genomic DNA (Promega). (E) Post-processed images from (D) where a binary output is obtained, white corresponds to amplification and black to non-amplification. (F) TTP values obtained from real-time fluorescence-based amplification using the *eluted disk* method from DBSFP and the elution from magnetic extraction with the Dynabeads kit.

Furthermore, extractions from whole blood stored in K_2_EDTA were evaluated with the *eluted disk* and *in-situ disk* methods independently (Figure 6B). Both methods showed a faster detection when the reverse transcriptase was incorporated in the reaction mix, demonstrating that both approaches are suitable for RNA extraction. However, a higher significant difference (p-value <0.0001) with and without the reverse transcriptase was observed with the *eluted disk* method.

### Colorimetric detection and comparison to a magnetic bead-based extraction method

The *eluted disk* method was compared against a commercially available magnetic bead-based nucleic acid extraction method (Dynabeads DNA DIRECT Blood, Invitrogen). For this comparison, LAMP colorimetric detection was chosen to evaluate the performance of the developed rapid extraction method from DBSFP from sample-to-result. For objective readout, an algorithm was implemented in MATLAB based on the work reported by *Rodriguez-Manzano et al*.^24^. The script was adjusted for the discrimination between pink and yellow to obtain a binary output where white corresponded to amplification and black to non-amplification. Workflow is shown in Figure 6C and script in Figure S5.

Results are shown in Figure 6D-E, where the amplification is visualised as a colour change from pink to yellow or as a binary output, respectively. Both methodologies, *eluted disk* from DBSFP and magnetic bead-based NAE from whole blood resulted in the amplification of all the samples and their objective readout with the software. As reference, real-time fluorescence-based detection was performed in parallel (Figure 6F). TTP values obtained in the real-time instrument were comparable between both methods (p-value > 0.05) in the presence of reverse transcriptase. Average TTP values of 5.99 ± 0.62 min and 6.34 ± 0.68 were obtained with *eluted disk* and magnetic bead-based NAE methods, respectively. Significant difference was observed with the *eluted disk* method with and without reverse transcriptase (5.99 ± 0.62 min and 16.12 ± 2.08, respectively). On the contrary, no significant difference was observed with the magnetic bead-based NAE method (6.34 ± 0.68 min and 6.52 ± 0.81, respectively). Lastly, the lower sample input for NAE from DBS (2 µL × 3 disks) compared to magnetic extraction (100 µL) demonstrates that this method is less invasive but equivalent in performance.

## Discussion

Molecular diagnostics from DBSFP envisioned to be used at the POC are currently limited by the complex laboratory-based methods for NAE^25,4^ or their application to PCR which require high temperature incubation and thermal-cycling.^7,15^ To date, a simplified and rapid protocol for NAE from filter papers and the subsequent downstream application for isothermal amplification has not been reported. In this work, we developed an electricity-free and rapid method for NAE from DBSFP (< 7 min) which allows RNA and DNA purification. Leveraging on the properties of DBSFP, two methods have been developed: *eluted disk* and *in-situ disk*.

The proposed NAE methods from DBSFP consisted of a pre-treatment of the filter paper with a mild surfactant (e.g. igepal^26–28^), spotting blood, a simple washing step (1 min) and incubation at RT (5 min). The elution obtained from the incubation step (*eluted disk*) and the use of the disk itself (*in-situ disk*) were successfully applied in combination with LAMP for the detection of the human reference gene ACTB. This is the first study that explored this duality of DBSFP. Typically, FTA classic cards are washed several times with proprietary FTA purification reagent and TE 1X buffer (∼10 min) and a drying step at 55 °C (∼15 min) is required prior to the use of the disks in molecular-based amplification; or in the case of FTA elute cards, several washing steps with proprietary FTA purification reagent (∼10 min) and incubation at 95 °C (∼30 min) is required before the elution is used for downstream molecular applications.^12,29^ In this work, we demonstrated that FTA classic cards, FTA elute cards and other filter papers (903 Protein Saver, Fusion 5 and Grade 3 filter paper) can be used with the *eluted disk* and *in-situ* disk methods for rapid and electricity-free NAE. Given the high cost of FTA cards, Fusion 5 or 903 Protein Saver (£3-5 per card) compared to grade 3 filter paper (∼£0.10 per card), we propose the use of the latter.

We also demonstrated RNA extraction from DBSFP without the need of equipment, long protocols or commercial kits^30–32^ as a proof-of-concept. Extraction with the *eluted disk* method with grade 3 filter paper from blood stored in K_2_EDTA showed significantly different results with and without the reverse transcriptase (6.99 ± 0.45 min and 18.76 ± 1.34). However, this was not the case for whole blood stored in LiHep. This anticoagulant may enhance the elution of DNA molecules such that the reverse transcriptase effect is negligible, which could be due to the negative charge of LiHep and its interaction with the DNA molecules in the matrix. Several studies have shown lower recovery of RNA from whole blood samples stored in LiHep compared to other anticoagulants such as K_2_EDTA.^33–35^ Nevertheless, further research is needed to fully understand the role of the anticoagulant in the preservation of RNA and DNA, since they could also be part of the pre-treatment of the DBSFP depending on the application and target. In future work, several applications in genomic and transcriptomic studies could be carried out involving the detection of endogenous genes by combining the *eluted disk* method from DBSFP with RNA-specific LAMP assays.

Finally, the *eluted disk* method showed comparable results to a magnetic bead-based extraction method using reverse transcriptase LAMP amplification and TTP values for comparative analysis. Average TTP values of 5.99 ± 0.62 min and 6.34 ± 0.68 were obtained with *eluted disk* and magnetic bead-based NAE methods, demonstrating that the *eluted disk* method from DBSFP is a potential alternative only requiring few microliters of blood. In future work, the applicability of this method could be extended to the detection of bloodstream pathogens such as virus, bacteria or parasites.^36–38^ The use of DBSFP without the need of prior cell or bacteria culture, will highly reduce the processing time of samples and allow the detection of pathogens which cannot be currently cultured. In addition, given the reported applicability of FTA cards for sample preservation and NAE, we expect that the proposed methods will be compatible with other fluid samples such as buccal swabs, faecal swabs, urine or saliva.^39^ Since we have also demonstrated a sample-to-result proof-of-concept workflow through the combination of the *eluted disk* method with colorimetric LAMP, we hope that both *elute disk* and *in-situ disk* could be used at the POC once embedded in a portable diagnostic format.^40^ Furthermore, we envision that once integrated they could be applied for sample archiving and improve the current pipeline for POC diagnostics where a massive number of tests are performed with loss of data.

## Author Contributions

All authors confirmed they have contributed to the intellectual content of this paper and have met the following 3 requirements: (a) significant contributions to the conception and design, acquisition of data, or analysis and interpretation of data; (b) drafting or revising the article for intellectual content; and (c) final approval of the published article. KMC and JRM conceived the idea and designed the study. KMC carried out the experiments and data analysis. KMC and JRM wrote the manuscript. JRM and PG supervised the project. JRM, PG, AC and JB proof-read the manuscript.

## Notes

### Competing Interest Statement

The authors have declared no competing interest.

